# Association of Vancomycin with Lipid Vesicles

**DOI:** 10.1101/160572

**Authors:** Xiaohui Hu, Kin Yip Tam

**Author notes:** **Corresponding author:** Prof. Kin Tam.

## Abstract

Antibiotics plays a pivotal role in modern medicine for the treatment of bacterial infection in patients. Membrane defines the boundary between single cell and its environment and is a main target for antibacterial agents. To better understand the mechanism of antibiotics action on microbes, we utilized liposome as membrane mimic model to study antibiotics interaction with bacterial membrane by variety of biophysical methods. Isothermal calorimetry and fluorescence photometry experiments were performed to examine interaction between antibiotics and liposome. We found that vancomycin, one of the most important antibiotics for the treatment of serious infections by gram-positive bacteria, binds to the liposome. The association between the drug and the liposome does not involve the tail part of the lipids. Moreover, the binding affinity increases along with the increment of liposome size. Of three major lipid components, phosphatidylglycerol is the preferential target for vancomycin binding. We also showed that vancomycin associates with vesicle derived from *Staphylococcus aureus* membrane in a similar manner as the binding to liposome. Our data suggested that vancomycin associates with bacterial membrane through direct interaction with lipid head groups with the extent of the association depending very much on specific type of lipids and curvature of local membrane structure.

## Introduction

Pathogenic bacteria pose a major threaten to public health, especially in under-developed region of the world. Large scale production and use of antibiotic penicillin during the World War II is a hallmark of modern antimicrobial therapy. Since then, a plethora of antimicrobial agents have been developed and commercialized, playing significant roles in improvement of human health, highlighted by gradual increasing of average life span worldwide. Prevailing of antibiotic treatment improves human health on one hand, but also triggers resistance of microbes to the drugs(Fischbach and Walsh 2009; Silver 2011).

Depending on their structure and target sites within microorganism, different antibacterial drugs have distinct modes of action(Kohanski et al. 2010). Of them, related to cell membrane are: inhibition or block of cell wall synthesis, inhibition of membrane protein or lipid synthesis and interference of cell membrane function. Some antibacterial peptides(Zasloff 2002; Brogden 2005), peptidomimics(Velkov et al. 2010) and small proteins(Mukherjee et al. 2014) could interact with or insert into cell membrane, where they interfere membrane formation or alter local membrane structure, thus reduce bacteria growth or reproduction. Bacterial membrane is surrounded by a layer of peptidoglycan cell wall. This unique feature had rendered great interest in searching for glycopeptide and lipoglycopeptide(Kahne et al. 2005), which mimics bacterial cell wall synthesis substrates or binds to synthesis intermediates, thus inhibits cell wall synthesis, while having little effect on host cells.

Glycopeptide antibiotic vancomycin has been used to treat a number of infect io ns caused by broad spectrum of gram-positive bacteria(Breukink and de Kruijff 2006; Stevens 2006). It inhibits cell wall formation by binding to D-Ala-D-Ala terminus of peptidoglycan precursors and preventing them from cross-linking(Wright and Walsh 1992; Eggert et al. 2001). To cope with rising of antibiotic resistance including vancomycin, developing of agents with multiple action modes had been a promising strategy(Otvos et al. 2005; Guilhelmelli et al. 2013; Yarlagadda et al. 2015). Herein, we studied the binding property of vancomycin to artificial and bacterial membranes using a variety of biophysics methods with the aim to obtain a better understanding of the binding process for expanding its antibiotic repository.

## Materials and Methods

### Materials

1-palmitoyl-2-oleoyl-*sn*-glycero-3-phospho-(1'-*rac*-glycerol) (sodium salt) (POPG), 1-hexadecanoyl-2-(9Z-octadecenoyl)-sn-glycero-3-phosphoethanolamine (POPE), 1-Oleoyl-2-[12-[(7-nitro-2-1,3-benzoxadiazol-4-yl)amino]dodecanoyl]-s n-Glycero-3-Phosphocholine (NBD PC) and cardiolipin (CL) were purchased from Avanti Polar Lipids. Vancomycin and nafcillin were from Sigma. Lysozyme and RNaseA were purchased from Merck. *Staphylococcus aureus* was from ATCC (strain #43300) and cultured in tryptic soy broth medium (Becton, Dickinson and Company). Other chemicals are obtained from Sigma if otherwise stated.

### Liposome preparation

Lipids of known mass in chloroform were well-mixed, then dried at room temperature in a glass tube by continuous purging of high pure nitrogen gas. Being thoroughly dried in a speed vacuum, the lipids were re-dissolved in PBS buffer pre-warmed over 45°C. After repeated pipetting, the hydrated lipids were applied to a Lipex Extruder (Northern Lipids Inc.) equipped with polycarbonate membrane for more than 10 times. Scattering profile of the liposome was then measured on a BI-200SM goniometer (Brookhaven Instruments) while being excited by a laser passing 637nm filter with 100 micron pinhole from 90 degree angle at 25°C. Data was collected and processed by the dynamic light scatter software coming with the equipment.

### Isolation of membrane from Staphylococcus aureus

Isolation of membrane were carried out as described(Poole 1993). Briefly, bacteria were cultured till late exponential phase before harvest. Cells was then re-suspended in 50 mM potassium phosphate buffer, pH8.0 and digested by lysozyme, deoxyribonuclease-1 and ribonuclease for 30 minutes in the presence of 10 mM MgSO_4_ at 37°C. The reaction was stopped abruptly by 15 mM EDTA for 1 minute. Additional MgSO_4_ was then added to the system to bring its final concentration back to 10 mM. Digested cell was centrifuged at room temperature, 25000g for 45 minutes to pellet the membrane, which was then re-suspended in PBS and applied through 18-g needle for more than 20 times. Resultant membrane vesicle was quantified by measuring protein concentration (Pierce^™^ BCA Protein Assay, Thermo Scientific), assuming equivalent amount of protein and lipid. Size distribution of the vesicles was assessed on a BI-200SM goniometer as describe in previous section.

### Isothermal calorimetry measurement

In a typical isothermal calorimetry(ITC) experiment carried out on a MicroCal iTC200 calorimeter (Malvern Instruments), 50 mM antibiotics were titrated into liposome with 5 mM lipid (total or individual) in PBS buffer at 25°C. After each injection, system was allowed to regain thermal balance between sample and reference chambers for 180 seconds with continuous stir of 750 rpm. Acquired data were processed by ITC-costumed Origin7 software using one-site model to avoid complication. Reference subtractions were performed against the data obtained by titrating antibiotic into PBS buffer.

### Fluorescence spectroscopy

Spectra of the fluorescent lipid incorporated liposome were collected on an LS55 Fluorescence spectrometer (PerkinElmer) at room temperature. The sample was excited by xenon lamp center at 485 nm with 2.5nm slit width. Antibiotic with high concentration was then introduced to minimize volume change. After mixed well, the spectrum of the sample was measured with the same setting.

## Results

### Vancomycin binds to liposome with a millimolar affinity

We first utilized liposome as membrane mimic to explore if vancomycin associates with the cell membrane at all. We performed isothermal calorimetry experiment by titrating antibiotics into liposome extruded through polycarbonate membrane (0.1 μm diameter). We found that vancomycin, but not β-lactam antibiotic nafcillin, introduced significant amount of heat change in liposome prepared from a mixture of POPE, POPG and cardiolipin (CL) (75:20:5) (Figure 1), a model mimic for bacterial membrane(Mary T. Le 2011). Fitting of the data into one-site model gave rise to millimolar level affinity (dissociate constant, *K*d), which indicated that vancomycin weakly, but definitely interact with liposome. The observation is in line with previous studies, where weak binding of vancomycin to model membrane were assessed by ITC(Al-Kaddah et al. 2010) and surface plasmon resonance(Kinouchi et al. 2014) techniques.

**Figure 1.**
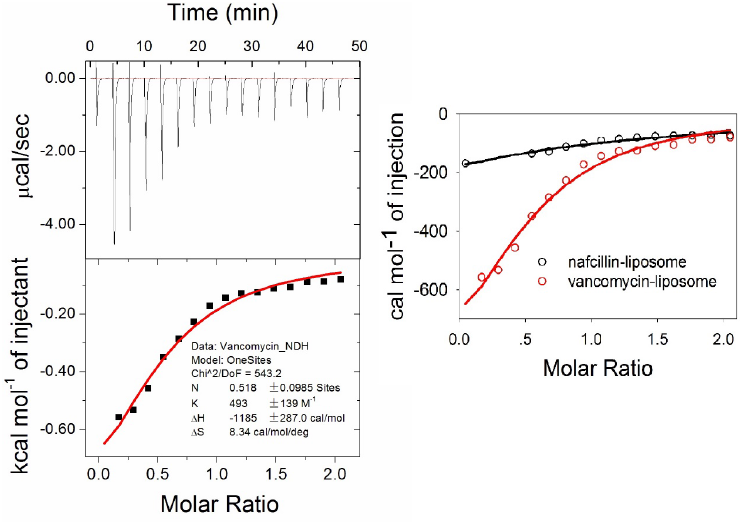
Titration of vancomycin into liposome. Raw data and fitting curve of ITC experiment (left); comparative fitting plot of naficillin and vancomycin titration with liposome (right).

### Vancomycin binds to vesicle derived from Staphylococcus aureus membrane

Now that we showed vancomycin association with membrane mimic liposome, it is tempting to ask whether the antibiotic can also bind to vesicle derived from biological sample. We thus isolated membrane from gram-positive bacterium *Staphylococcus aureus* by means of enzymatic digestion, mechanical disruption and syringe needle homogenization(Poole 1993). Resultant vesicle was then undergone calorimetric titration with vancomycin as the titrant. According to the fitting result of ITC data, vancomycin also bound to the bacterial derived vesicle (Figure 2.A), but with much higher affinity (sub-millimolar) than that of liposome (Figure 2.B). Although we cannot totally rule out the possibility of residual peptidoglycan remaining on the membrane, lysozyme digestion step of the cell should remove most, if not all of them. Apparently, vancomycin binds to the bacterial membrane derived vesicle with higher affinity.

**Figure 2.**
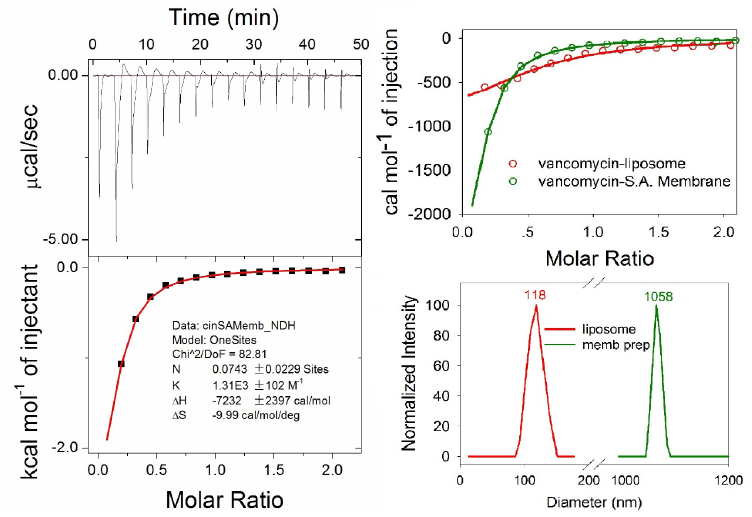
Titration of vancomycin into vesicle derived from *Staphylococcus aureus* membrane. Raw data and fitting curve of ITC experiment (left); comparative fitting plot of vancomycin titration with liposome and vesicle from membrane (top right); size distribution of liposome and membrane vesicle determined by dynamic light scattering (bottom right).

### Vancomycin strengthens the association with liposome as its size increases

In light of differential binding between liposome and membrane-derived vesicle to vancomycin and the fact that the later measured much larger in diameter than the former (Figure 2), we speculated that the size of liposome/vesicle might play a role in the antibiotic association with membrane mimic. To test the hypothesis, we performed serial calorimetric titration of vancomycin into liposome prepared with polycarbonate membranes of various size (0.1, 0.4 and 1 μm). We found that the binding affinity of vancomycin increased along with liposome size increment. (Figure 3), which indicated that cell surface landscape or/and membrane curvature could be an important factor in vancomycin association.

**Figure 3.**
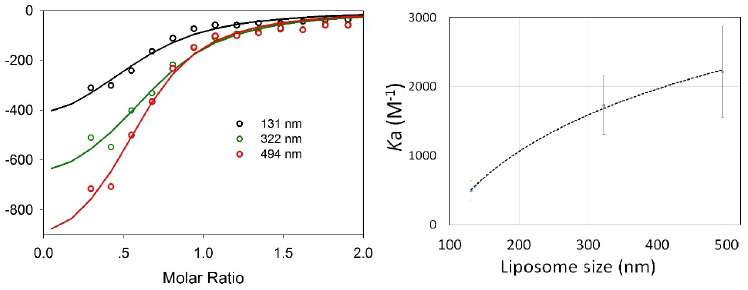
Size-dependent binding of liposome with vancomycin. Data and fitting curve from titrations of liposome were plotted in same figure for comparison (left); *K*a value derived from fitting were plotted against the sizes of the liposome (right).

### Vancomycin binds preferentially to POPG over cardiolipin and POPE

Because the lipids used in the study are similar structurally in fatty acid moiety but vary in head groups. We reasoned that their difference in head group could contribute differently to the association with vancomycin. To explore binding preference of vancomycin among lipids, we performed ITC measurement of the antibiotic with liposomes prepared from individual lipid. Based on the *K*a value derived from data fitting, binding affinity of vancomycin to POPG is significantly higher than that to POPE and CL (Figure 4). Thus, we established that POPG is the major contributor to the association of vancomycin with liposome The observation could be explained by careful examination of the structure of vancomycin(Loll et al. 1997; Jia et al. 2013) and the lipids (Figure 4). Positive charged amine in the head group of POPE does not appear to favor the interaction with uncharged vancomycin. Moreover, embedded hydroxyl group on hydrocarbon chain within two phosphoryl group is unlikely to facilitate binding in the case of cardiolipin. On the other hand, the two hydroxyls on the head group of the POPG could potentially interact with vancomycin via hydrogen bond formation, which promotes the glycopeptide association with lipid membrane(Domenech et al. 2009).

**Figure 4.**
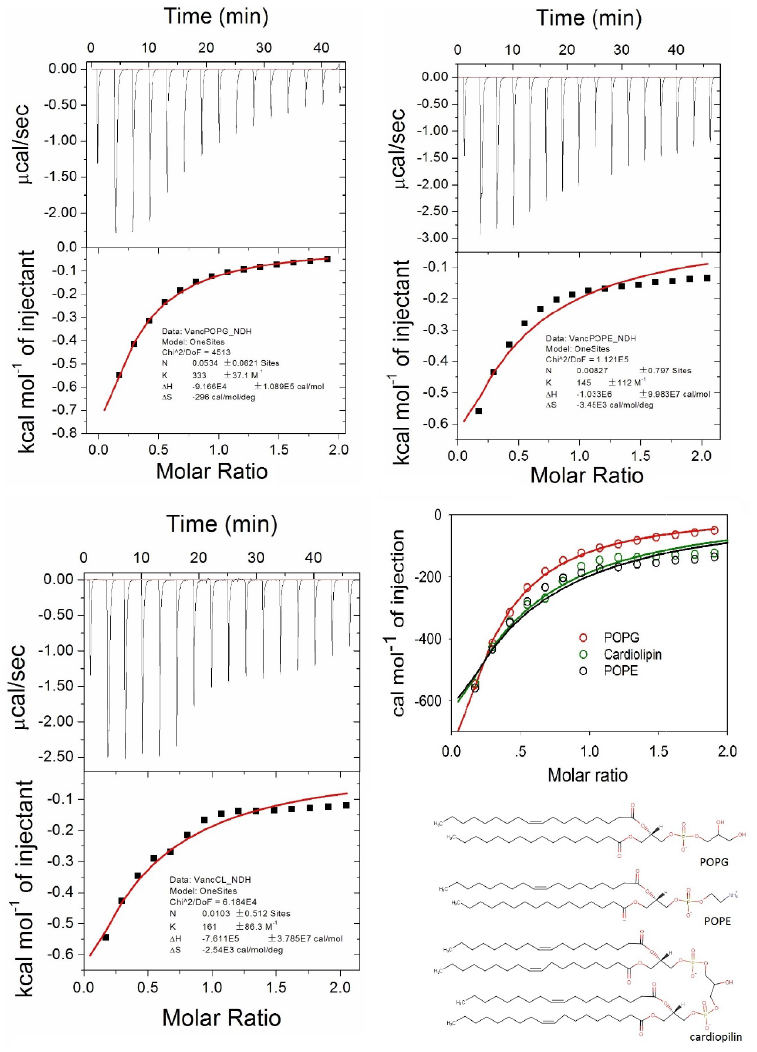
Vancomycin binds preferentially to POPG over POPE and cardiolipin. ITC raw data and fitting of vancomycin titration with POPG (top left), POPE (top right) and cardiolipin (bottom left); Fitting of titrations were plotted for comparison (middle left); Structure of three lipids were shown (bottom right).

### Vancomycin interaction with liposome does not involve acyl terminus of the lipids

We have shown that vancomycin associates with liposome through its interaction with the head group of the lipids. But it remains unclear if the antibiotic could introduce any change inside the lipid bilayer. Fluorescence spectroscopy was performed to gain more insight into the binding of vancomycin to liposome. We incorporated trace amount (0.5%) of tail fluorescence-labeled lipid (NDB-PC) into liposome. Addition of stoichiometric amount of vancomycin did not induce significant change in the spectrum of the liposome (Figure 5), suggesting that binding of vancomycin to liposome might not involve the acyl terminus of the lipid. In other words, it is unlikely for the antibiotic to insert deep into the lipids bilayer. Otherwise, alteration in fluorescence intens it y or/and peak position shift would be obvious because of fluorophore disturbing by the insertion, as seen in a control experiment, where pore formation depsipeptide antibiotics valinomycin(Andreoli 1967; Bhattacharyya et al. 1971) reduced liposome fluorescence intensity remarkably. This is consistent with the fact that, vancomycin and other glycopeptides generally exhibit low membrane permeability, which has been well documented in literatures(Domenech et al. 2009; Kohanski et al. 2010).

**Figure 5.**
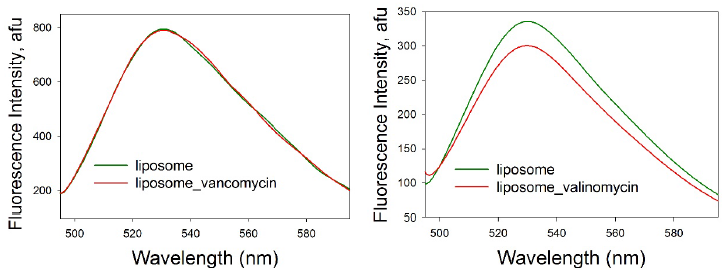
Fluorescence spectra of liposome with fluorescent label. Left panel, spectra of 3.5 mM liposome containing 0.5% NDB PC in the absence (black line) and presence (red line) of 5 mM vancomycin; right panel, spectra of 1.5 mM liposome containing 0.5% NDB PC in the absence (black line) and presence (red line) of 2 mM valinomycin.

## Discussion

We have shown that glycopeptide antibiotics vancomycin associate with biomimetic membrane, a fact have been largely ignored in the field because of its weak interaction(Al-Kaddah et al. 2010; Kinouchi et al. 2014). Furthermore, we showed for the first time that it binds to membrane vesicle derived from gram-positive bacteria *Staphylococcus aureus* in a similar fashion. These observations could potentially open a new avenue for the prototype antibiotics to tackle infectious microbes by blocking cell wall synthesis and interfering cell membrane function simultaneously. Multip le mechanism of action is thought to be an effective strategy to combat with prevalent resistance nowadays rising among pathological microorganism.(Guilhelmelli et al. 2013; Gardete and Tomasz 2014; Yarlagadda et al. 2015)

We also found that vancomycin appears to bind to the membrane from outside and the extent of association depends much on local phospholipid composition and the curvature of the membrane. POPG appears to be the favorite target for vancomycin binding and larger size (flattened surface) of the membrane helps too. These finding could provide a framework for further modification of the antibiotics to optimize its binding to plasma membrane and thus higher potency. To this end, a further investigation of vancomycin structure and bacterial membrane composition is required. In addition, dimerization of vancomycin(Loll et al. 1997; Jia et al. 2013) might also play an important role as a larger hydrophobic interface once shielded by dimeric interaction has to be covered by membrane association.

## Conclusions

We have studied the binding of vancomycin to biomimetic membranes based on three phospholipid components and bacterial membranes by using a variety of biophysic a l methods, including isothermal calorimetry and fluorescence photometry. It has been shown that the ways of association of vancomycin to biomimetic membranes is very similar to that of vesicle derived from *Staphylococcus aureus* membrane. Our results indicated that vancomycin associates to the cell membrane from outside, and does not permeate through the cell membrane, which is consistent with the mechanisms of the drug action. By varying the size and composition of the biomimetic membrane vesicles, it has been demonstrated that the extent of association depends on specific type of lipids and curvature of local membrane structure. These observations could potentially open a new avenue to revive the prototype antibiotic to fight infectious microbe through the design of specific tight binders to the outer surface of bacterial membranes to impede cell wall synthesis.

## Acknowledgement

We thank the financial support from the University of Macau (grant no.: SRG2013-00055-FHS). We thank Prof Jun Zheng (University of Macau) for providing a bacteria strain of *Staphylococcus aureus.*

